# Structure-based humanization of a therapeutic antibody for Multiple Myeloma

**DOI:** 10.1101/2023.09.27.559758

**Authors:** Stephen F. Marino, Oliver Daumke

## Abstract

The optimal efficacy of xenogeneically generated proteins intended for application in humans requires that their own antigenicity be minimized. This necessary adaptation of antibodies to a humanized version poses challenges since modifications even distant from the binding sites can greatly influence antigen recognition and this is the primary feature that must be maintained during all modifications. Current strategies often rely on grafting and/or randomization/selection to arrive at a humanized variant retaining the binding properties of the original molecule. However, in terms of speed and efficiency, rationally directed approaches can be superior, provided the requisite structural information is available. We present here a humanization procedure based on the high-resolution X-ray structure of a chimeric IgG against a marker for Multiple Myeloma. Based on *in silico* modelling of humanizing amino acid substitutions identified from sequence alignments, we devised a straightforward cloning procedure to rapidly evaluate the proposed sequence changes. Careful inspection of the structure allowed identification of a potentially problematic amino acid change that indeed disrupted antigen binding. Subsequent optimization of the antigen binding loop sequences resulted in substantial recovery of binding affinity lost in the completely humanized antibody. X-ray structures of the humanized and optimized variants demonstrate that the antigen binding mode is preserved, with surprisingly few direct contacts to antibody atoms. These results underline the importance of structural information for the efficient optimization of protein therapeutics.

## Introduction

The focus on antibodies as therapeutics continues to increase in parallel with clinical successes^1^. Their evolutionarily optimized functionality, above all their specificity, make them promising options for treating many human pathologies. Antibodies intended for therapeutic use are most often generated in non-human mammals, and therefore require some modification to minimize or eliminate their own antigenicity. Although the necessary mutations are identified by comparing the homologous heavy and light chain variable region (V_H_ and V_L_) subtypes from the source animal and humans, their introduction often leads to changes in the antigen binding properties even when the complementarity determining region (CDR) loops, i.e. the antigen binding sites, are not directly modified^2^. For this reason, several strategies intended to preserve binding have been developed. The first humanization attempts were made by grafting mouse variable regions onto human antibody frameworks^3-5^. Guided by crystallographic structures, improvements have been made by grafting only CDRs^6^ or only the SDRs (specificity determining residues, the residues known or assumed to directly interact with the antigen)^7^. In most cases though, negative effects on binding affinity result. Randomization/selection routines using phage display technology have been applied to address this problem by applying repeated cycles of variable chain replacement and subsequent selection for antigen binding, with reasonable success ^8-9^. In terms of the improvements gained vs. time/resources spent, it is nevertheless preferable to work in a directed fashion, for which structural information is a prerequisite.

Multiple Myeloma (MM) is a currently incurable malignancy of plasma cells (PCs) and accounts for nearly 1% of all cancer occurrences^10^. It is the second most frequently diagnosed blood cancer with 150,000 new cases globally per year and a rising incidence – likely due to increasing lifespans and the fact that most cases are diagnosed in patients above age 65^11^. The median survival for MM patients has been extended in recent years, due primarily to the use of proteasome inhibitors in combination with immune modulators and thalidomide analogs^12^. However, because of the near ubiquitous eventual resistance to these treatments, antigen targeted approaches have become a main focus in the development of new therapies for MM.

One of the most attractive targets for both antibody and Chimeric Antigen Receptor T-cell (CAR-T) based treatments is the B-cell Maturation Antigen (BCMA). This cell surface receptor is involved in the survival of long lived PCs in the bone marrow via interactions with its native soluble ligands APRIL and BAFF^13-15^. BCMA is exclusively present in high copy number on long lived PCs (and in only very low copy on some other types of malignant B-cells^16^) and this specificity makes it a highly desirable target for MM treatment. Although a number of monoclonal IgGs have been generated against BCMA, only one has been thoroughly structurally characterized, the chimeric mouse/human J22.9-xi that showed substantial anti-tumor effects in an MM mouse model^17^. The high resolution X-ray structure of J22.9-xi in complex with the extracellular domain of BCMA provided fine details of the binding interaction and the only completely verified BCMA epitope, which overlaps those of both native BCMA ligands.

Having shown promise in retarding tumor growth in mice, it was desirable to generate fully humanized J22.9-xi variants for eventual testing in MM patients. Using the structure as a guide, we devised a mutagenesis scheme allowing rapid generation and testing of humanized antibodies retaining high affinity binding and full functionality. We describe here the procedure and provide further structural information on both humanized and CDR optimized variants, verifying the retention of the antigen binding mode despite extensive mutagenesis. The success of this rapid humanization procedure emphasizes the importance of structural information for medical advances.

## Materials and Methods

### Protein production and purification

The production, purification and generation of Fab:BCMA complexes and crystallization studies were all performed as previously described^17^.

### Humanization of J22.9

The residue alterations required to produce fully humanized sequence variants were chosen based on sequence alignments to the human heavy and light chain variable region sequences corresponding to those of the mouse J22.9-xi. Each modification was first assessed *in silico* by modelling them into the crystal structure of the J22.9-xi:hBCMA complex using Coot^18^. After flagging potentially problematic mutations, two complete J22.9 variable region genes for each chain were synthesized, one of the original mouse sequence and one humanized sequence containing the complete set of humanizing mutations. The nucleotide sequences were designed to introduce two unique restriction enzyme sites, dividing the genes into three cassettes each. Various combinations of the original mouse and fully humanized gene cassettes were produced, the resulting IgGs expressed, purified and subjected to qualitative screening via FACS using BCMA-positive cells (as described in^17^). Problematic residues and residues for CDR optimization of the final humanized antibody were individually replaced via PCR as needed to regain/improve antigen binding. The final constructs were then quantitatively assessed for binding to both human and cynomolgus BCMA (hBCMA and cBCMA, respectively) via SPR.

### Surface Plasmon Resonance (SPR)

Affinity measurements by SPR were performed on a ProteonXPR36 in phosphate buffered saline with 0.005% Tween-20 (PBST). The respective, whole IgGs (15 ug/ml) were immobilized on a Proteon GLH sensor chip using standard amine chemistry according to the manufacturer’s instructions. PBST containing human or cynomolgus BCMA served as the mobile phase. The binding affinity (Kd) was calculated from association (k_on_) and dissociation (k_0ff_) constants determined in parallel on a dilution series of BCMA (ranging from 0.4 to 800 nM for hBCMA and 2.7 nM to 1 μM for cBCMA) assuming a single-site binding model.

### Crystallography

All crystals were grown using Cu(II) containing screens developed for the original J22.9-xi Fab fragment:hBCMA complex, comprising 21% PEG 3350, 0.1 M BisTris pH 6.5 and 5 mM CuCl_2_ at 20° C. Crystals were screened to assess diffraction quality using an HC-1 dehydration device^19^ on Beamline 14.3 of the BESSY II Synchrotron at the Helmholtz Zentrum Berlin (HZB), Germany. Individual crystals were extracted from drops using meshes and mounted on a goniometer under the HC-1 air stream at 99% humidity at room temperature. Mother liquor was removed from around each crystal manually using fine paper wicks, leaving the naked crystals in direct contact with the humidified air stream after which diffraction images were taken and evaluated. Crystals intended for data collection were then directly frozen without cryoprotectant by rapidly covering them with cryovials filled with liquid nitrogen and immediate removal from the goniometer to a liquid nitrogen bath. Frozen crystals were stored in liquid nitrogen until data collection at Beamlines 14.2 and 14.3 of the HZB. Structures were solved by molecular replacement with Phaser^20^ (J22.9-H using J22.9-xi as the search model and both J22.9-FNY and J22.9-ISY using J22.9-H). The structures were iteratively refined in Phenix^21^ with model adjustment and superpositions in Coot^22^ and PyMOL^23^. Figures were generated with PyMOL.

## Results

### Humanization of J22.9-xi

The variable domains from the original J22.9 mouse IgG had been cloned onto a human IgG_1_ F_c_ domain for recombinant production, generating the chimeric J22.9 -xi^17^. For the full humanization, identification of the human germline sequences corresponding to the J22.9-xi V_H_ and V_L_ chains was performed using IgBLAST^24^ and subsequent sequence alignments indicated a total of 41 residue changes needed to produce fully humanized sequences, 16 in V_H_ and 25 in V_L_ (Fig 1A, B). Each modification was modelled into the J22.9-xi:BCMA complex structure to identify any potential clashes or unfavorable bond angles due to steric constraints. Nearly all substitutions (mostly surface exposed) appeared likely to be accommodated by the antibody without obvious impact on the binding site.

**Figure 1.**
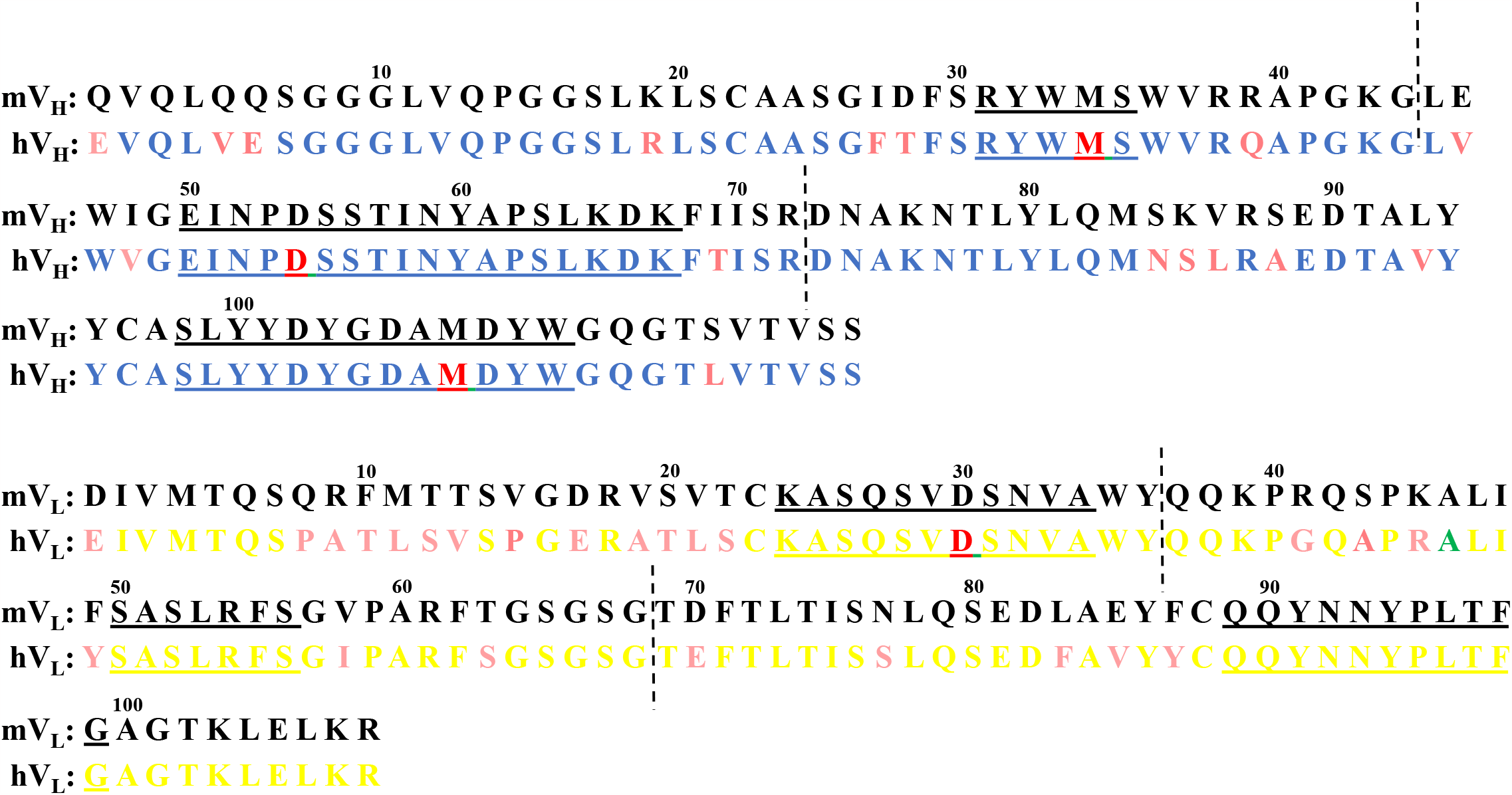
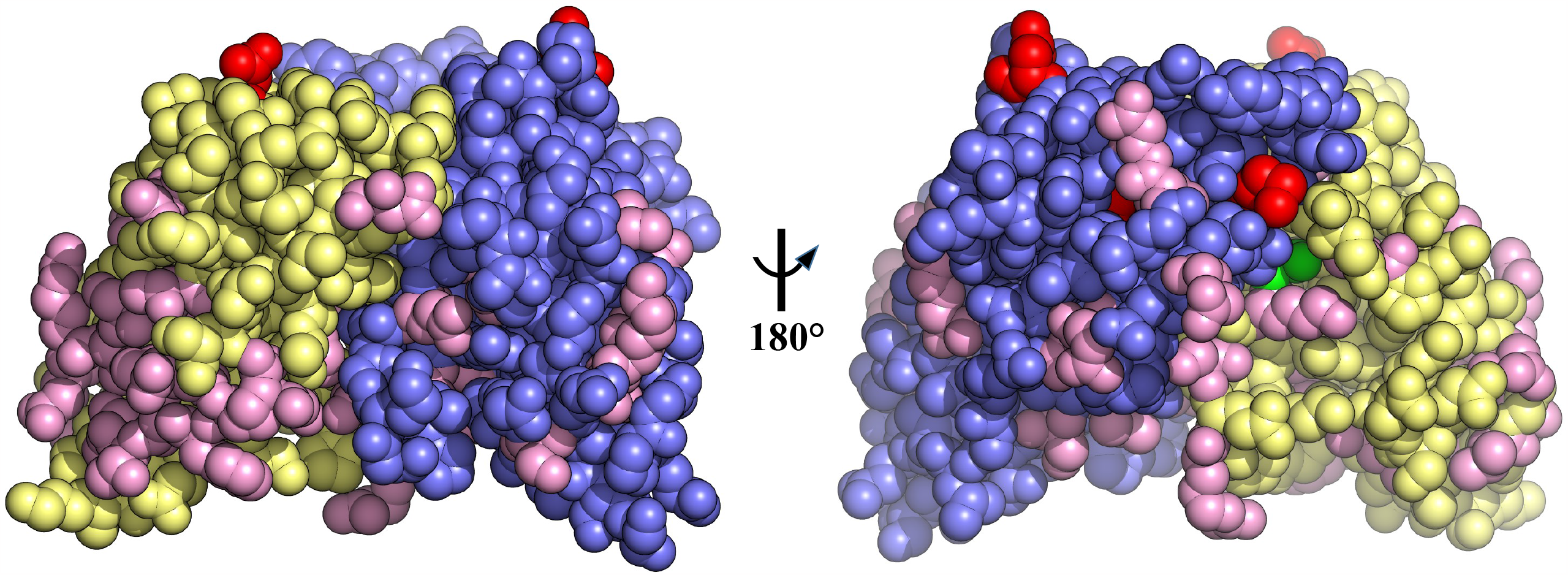
Humanizing mutations in J22.9-xi. A. Protein sequences of the mouse variable region heavy (mV_H_) and light (mV_L_) chains aligned with the corresponding human germline sequences (hV_H_ and hV_L_). CDRs 1, 2 and 3 from each chain are underlined. Humanizing mutations are highlighted in magenta, proposed CDR positions for stabilizing PTM mutations are highlighted in red and the complex disrupting A46 indicated in green. Vertical dashed lines indicate the corresponding boundaries of the cloning cassettes. B. Space-filling model of the J22.9-xi variable domain showing the positions of the proposed mutations on the structure (PDB ID: 4ZFO). As in (A), the V_H_ is depicted in blue, the V_L_ in yellow, humanizing mutation positions in magenta, stabilizing CDR PTM positions in red and A46 in green. The right panel view depicts a 180° rotation along the vertical axis of the left panel view.

The single exception concerned alanine 46 (A46) of the mouse V_L_, that in the human V_L_ is a leucine (variation = A46L). In the J22.9-xi:BCMA structure, A46 is buried in a hydrophobic pocket bounded by L99, D108 and W110 from V_H_ and Y36, L47 and F55 from V_L_ (Fig 2). These residues, together with the β-carbon from the side chain of D108 from V_H_, form a small cavity closely packed around the alanine β-carbon atom that is directly adjacent to the BCMA binding site. The sidechain faces of both F55 and L99 opposite A46 are in direct contact with L17 in BCMA, the residue at the center of the D*x*L loop and the primary recognition feature for J22.9-xi and both native BCMA ligands. Additionally, F49 from V_L_ is packed tightly against F55 from V_L_ and L18 of BCMA. The additional volume required upon substitution with the much larger leucine sidechain was expected to induce substantial rearrangement of the interactions around the A46 pocket and therefore interfere with BCMA binding.

**Figure 2.**
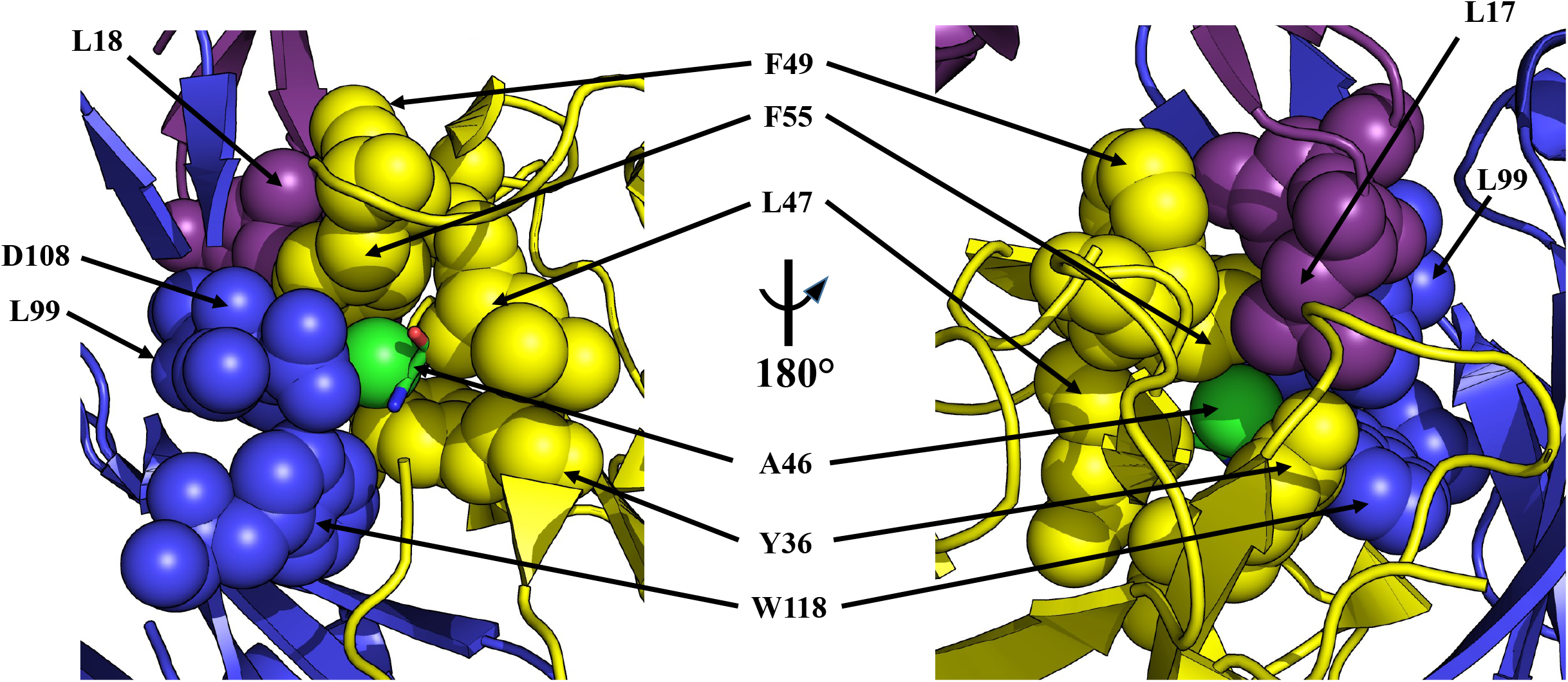
The A46 hydrophobic pocket of J22.9-xi. Two views of the A46 binding environment with corresponding residues in space-filling representation and labeled. V_H_ is in blue, V_L_ in yellow and BCMA in violet. A46 (V_L_) is seen in the center in stick representation with color-coding of individual atoms (nitrogen blue, carbon green and oxygen red) and the beta carbon atom in the pocket represented as a green sphere. The right panel view depicts a 180° rotation along the vertical axis of the left panel view. The close packing around the A46 sidechain is clear from both panels as are the interactions with L17 and L18 of BCMA.

We divided the positions to be mutated between cloning cassettes that could be easily combined to test the maximum number of substitutions with as few new constructs as possible (Fig. 1A). It was reasoned that by stepwise combination of cassettes encoding the fully humanized sequences with those of the original mouse sequences, detrimental mutations (as qualitatively assessed by binding to cells displaying BCMA in FACS, described in^17^) could be rapidly identified. Mutated positions on cassettes whose presence disrupted binding could then be individually replaced by PCR to narrow down the problematic change(s). Several constructs were cloned and the antibodies produced in small scale as described for J22.9-xi. It was quickly determined that only antibodies harboring the A46L substitution were negative for binding by FACS (not shown). Reversing the change back to alanine restored binding in the absence of any other changes, confirming the original predictions from the modelling. The final humanized variant comprising all substitutions in both the V_H_ and V_L_, with the exception of A46 in the V_L_, is hereafter referred to as J22.9-H.

### CDR stability optimization

The affinity of J22.9-H for human BCMA (hBCMA) was determined by SPR to be 1.5±0.3 x 10^-8^ M – a nearly 200-fold loss compared to J22.9-xi at 2.8±0.7 x 10^-10^ M (Table I and Figure 3). In an attempt to mitigate this loss in affinity and impart greater long-term stability to the variable domains, we modelled additional mutations into the CDRs of both chains that were intended to prevent detrimental post-translational modifications (PTMs). The sulfur containing methionine is subject to oxidation by reactive oxygen species, particularly at low pH, and this has been shown to reduce the conformational and thermal stability of antibodies^25, 26^. In their ionized form, Asp sidechains can perform a peptide chain breaking reaction in which the sidechain carboxylate attacks the peptide bond carbonyl carbon^27^ and Asp residues adjacent to those having hydroxyl containing sidechains, like serine, have been shown to be more reactive^28^. Three proposed PTM sites were identified in hV_H_, one in each of the three CDRs (Fig 1A): methionine 34 in CDR1 to be substituted with isoleucine and phenylalanine (M34I/F); aspartic acid 54 in CDR2 to be substituted with serine (D54S); and methionine 107 in CDR3 to be substituted with tyrosine (M107Y). Only one mutation in hV_L_, aspartic acid 30 to glutamic acid (D30E) in CDR1, was proposed. Since both J22.9-H aspartic acid residues, D54 and D30, occur adjacent to a serine, both were targeted for replacement. Although conformational changes to the CDRs induced by the proposed substitutions could not be ruled out, inspection of the J22.9-xi:BCMA complex structure made clear that none of these residues made either direct or indirect (over water molecules) contacts to BCMA^29^. The final constructs incorporating all of the humanizing mutations (except A46L) and the stabilizing CDR PTM substitutions, 44 residue changes in total, are referred to as J22.9-FSY and J22.9-ISY, based on the identity of the hV_H_ M34 substitution (phenylalanine, F, and isoleucine, I, respectively).

**Figure 3.**
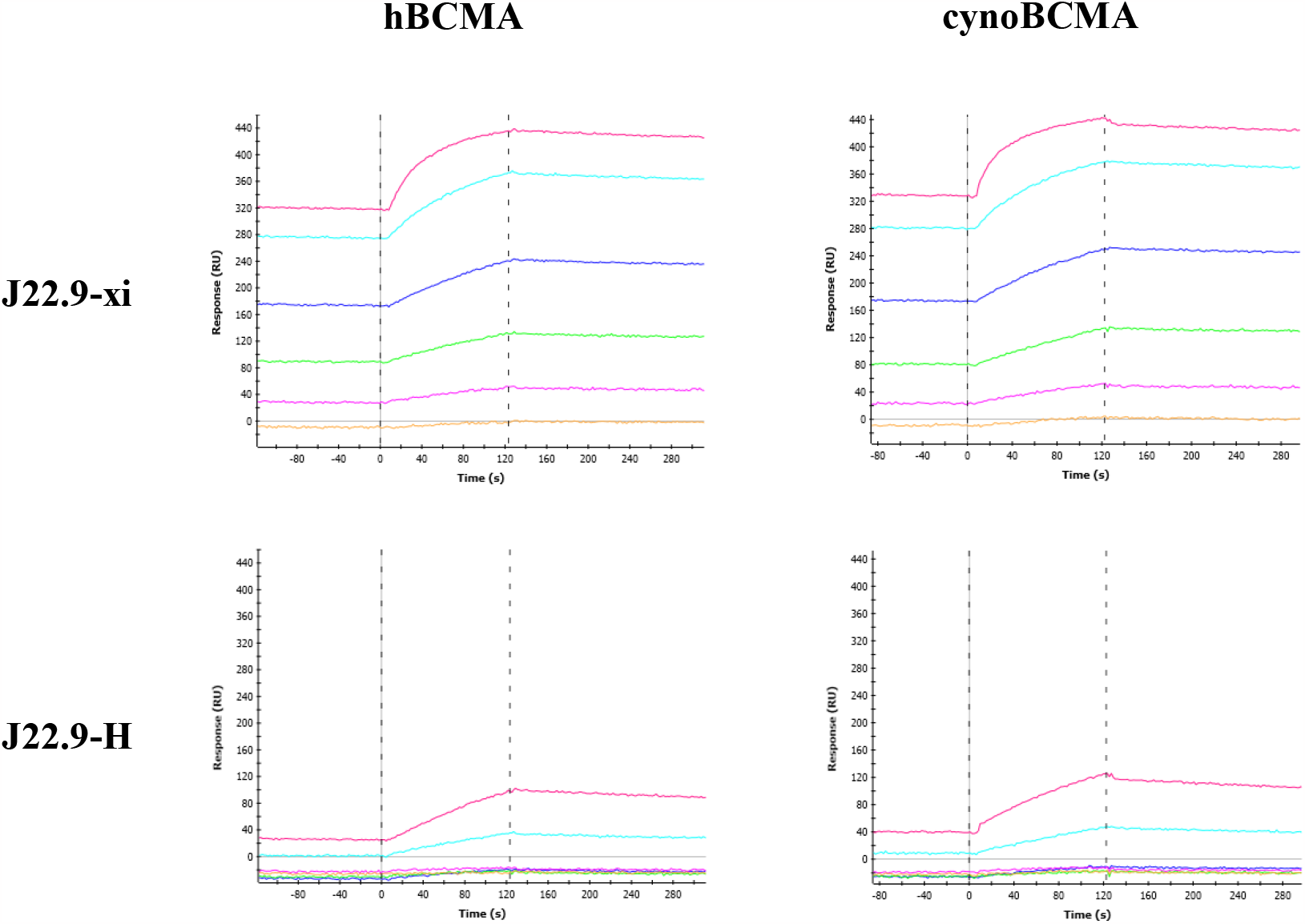
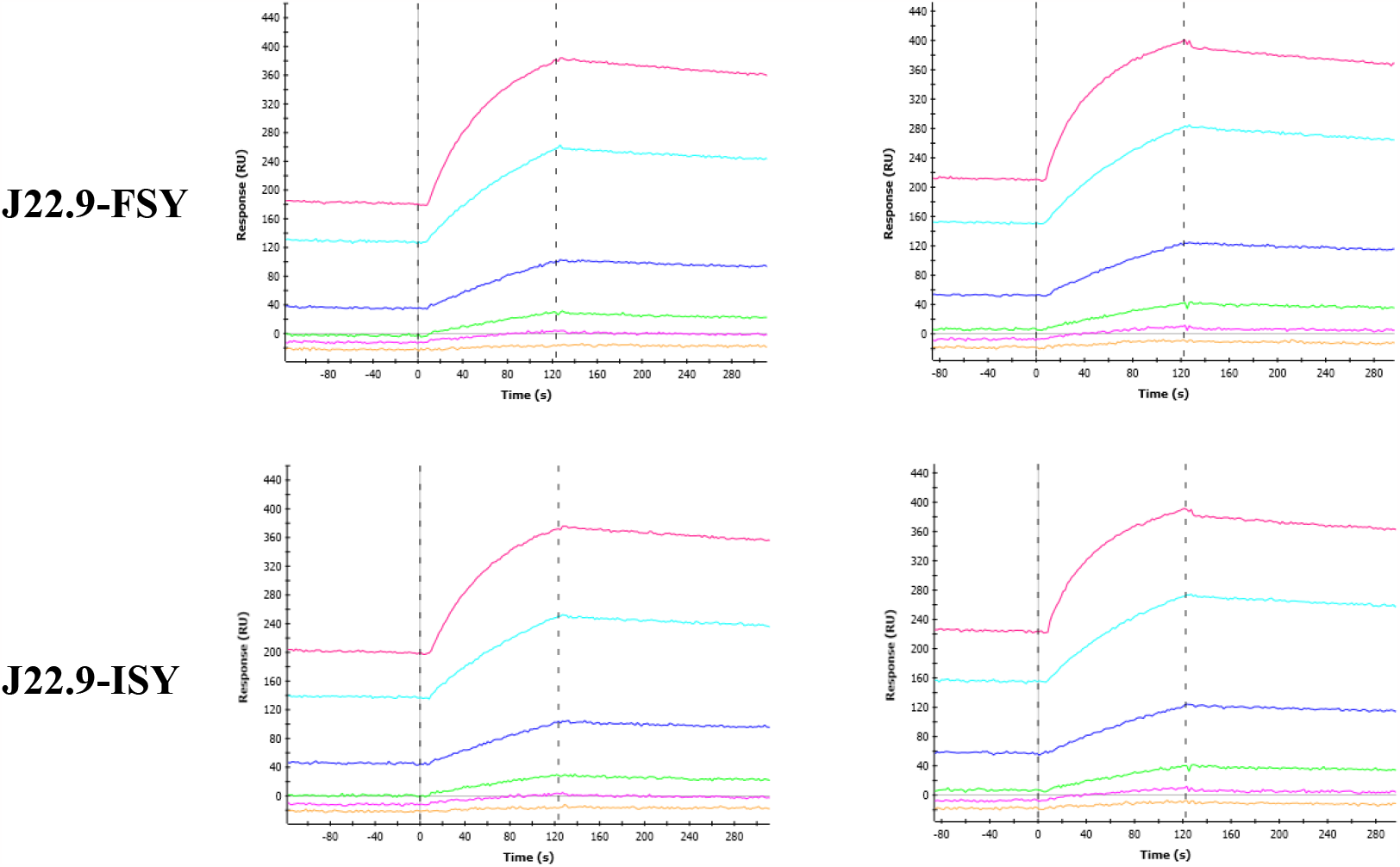
Measurement of BCMA binding affinities of J22.9 variants. SPR sensorgrams of J22.9 variants (labeled) binding to hBCMA and cynoBCMA. The colored traces correspond to increasing concentrations of BCMA in the mobile phase binding to immobilized IgG. The leftmost vertical dashed line indicates the start of ligand flow over the immobilized antibodies, during which association is measured, and the rightmost indicates the switch to BCMA-free buffer, initiating the dissociation phase.

**Table 1:** Affinities of J22.9 constructs for BCMA homologues.

| Construct | Affinity (SPR)<br>(human)<br>(n=3) | Affinity (SPR)<br>(cynomolgus)<br>(n=2) |
| --- | --- | --- |
| J22.9-xi | $2.8 \pm 0.7 \times 10^{-10} \text{ M}$ | $2.7 \times 10^{-9} \text{ M}$ |
| J22.9-H | $1.5 \pm 0.3 \times 10^{-8} \text{ M}$ | $2.0 \times 10^{-7} \text{ M}$ |
| J22.9-FSY | $2.2 \pm 0.3 \times 10^{-9} \text{ M}$ | $2.0 \times 10^{-8} \text{ M}$ |
| J22.9-ISY | $2.0 \pm 0.2 \times 10^{-9} \text{ M}$ | $1.7 \times 10^{-8} \text{ M}$ |

By SPR, the set of stabilizing CDR substitutions did indeed improve the binding affinity for BCMA, fortuitously by more than 10-fold over J22.9-H, putting them in the low nanomolar range (Table I). Since toxicity studies carried out in preparation for clinical trials often use macaques (cynomolgus monkeys), the affinity of each variant for cynomolgus BCMA (cynoBCMA) was also determined. cynoBCMA differs from the human homolog at three positions – in cynoBCMA, alanine 20 is an aspartic acid and isoleucine 22 is a lysine - a potentially destabilizing change for the interaction with the antibodies, since it introduces a charged side chain directly into the hBCMA binding epitope; additionally, asparagine 31 in hBCMA is deleted in cynoBCMA. However, despite the presence of lysine 22, all variants were able to bind cynoBCMA, and in all cases the affinity by SPR was approximately 10-fold lower for cynoBCMA than for hBCMA.

### Crystal structures

All variants were subjected to crystallization trials and diffracting crystals were obtained for J22.9-H and J22.9-ISY. We noted that a D54N substitution in the hV_H_ CDR2 would produce a eukaryotic N-linked glycosylation signal sequence (consensus sequence N–X–S/T, where X is any amino acid except proline; in J22.9-H, the D54N substitution produces N-S-S) and, since the CDR2 loop does not extensively contact BCMA, we introduced the D54N substitution into J22.9-FSY. With the resulting variant, referred to as J22.9-FNY, we also obtained diffracting crystals. We solved all three structures by molecular replacement: J22.9-H and J22.9-ISY to Å, and J22.9-FNY to 2.7 Å (Table II), thereby allowing comparison of the binding interaction of the fully humanized and optimized IgGs with that of the original chimeric and biologically active J22.9-xi.

**Table 2.** Data collection and refinement statistics.

|  | **J22.9-H** | **J22.9-FNY** | **J22.9-ISY** |
| --- | --- | --- | --- |
| **Wavelength** | 0.9184 | 0.9184 | 0.9184 |
| **Resolution range** | 34.78 – 3.1<br>(3.211 – 3.1) | 21.76 – 2.72<br>(2.817 – 2.72) | 45.25 – 3.094<br>(3.205 – 3.094) |
| **Space group** | C 1 2 1 | C 1 2 1 | C 1 2 1 |
| **Unit cell** | 137.36 55.18 81.16<br>90 121.023 90 | 137.266 55.371 69.486<br>90 108.009 90 | 137.185 54.617 81.084 90<br>121.089 90 |
| **Total reflections** | 39770 | 48615 | 34143 |
| **Unique reflections** | 9584 (942) | 13486 (1323) | 9514 (925) |
| **Multiplicity** | 4.14 | 3.60 | 3.59 |
| **Completeness (%)** | 99.19 (99.37) | 99.42 (99.55) | 98.96 (95.65) |
| **Mean I/sigma(I)** | 6.54 | 9.98 (2.03) | 5.15 (1.74) |
| **Wilson B-factor** | 68.29 | 49.68 | 63.36 |
| **Reflections used in refinement** | 9580 (941) | 13483 (1322) | 9502 (924) |
| **Reflections used For R-free** | 480 (47) | 675 (66) | 476 (46) |
| **R-work** | 0.2609 (0.4045) | 0.2440 (0.4010) | 0.2466 (0.3737) |
| **R-free** | 0.3109 (0.4196) | 0.2967 (0.3842) | 0.3056 (0.4203) |
| **Number of non-hydrogen atoms** | 3566 | 3597 | 3525 |
| **macromolecules** | 3521 | 3547 | 3524 |
| **ligands** | 2 | 30 | 1 |
| **solvent** | 43 | 20 |  |
| **Protein residues** | 461 | 464 | 461 |
| **RMS (bonds)** | 0.002 | 0.002 | 0.002 |
| **RMS (angles)** | 0.44 | 0.45 | 0.49 |
| **Ramachandran**<br>**favoured (%)** | 91.17 | 92.76 | 90.29 |
| **Ramachandran**<br>**allowed (%)** | 8.17 | 6.58 | 7.73 |
| **Ramachandran**<br>**outliers (%)** | 0.66 | 0.66 | 1.99 |
| **Rotamer**<br>**outliers (%)** | 0.76 | 0.75 | 1.26 |
| **Clashscore** | 2.45 | 2.84 | 3.03 |
| **Average B-factor** | 80.15 | 55.31 | 74.73 |
| **macromolecules** | 80.48 | 55.15 | 74.75 |
| **ligands** | 102.33 | 85.14 | 22.51 |
| **solvent** | 51.90 | 38.36 |  |
| **Number of TLS**<br>**groups** | 3 | 3 | 3 |

Statistics for the highest-resolution shell are shown in parentheses.

Alignment of J22.9-H and the fully humanized and CDR optimized J22.9-ISY and J22.9-FNY structures with that of J22.9-xi provides verification of the retention of the BCMA binding mode after incorporation of the necessary 44 substitutions (Fig. 4). The backbone root mean square deviation (RMSD) of the variable domains between J22.9-xi and J22.9-H is 0.88 Å, indicating no substantial changes in overall architecture. The V_H_ and V_L_ domain conformations of both J22.9-FNY and J22.9-ISY are, within experimental error, identical to those of the corresponding J22.9-H domains, demonstrating the minimal structural impact of the PTM modifications. There are only minor alterations in the backbone conformations of CDRs 1 and 2 from V_L_ between J22.9-xi and the humanized variants, while differences between the V_H_ chains are also minimal, confined to small shifts without changes in their topologies. Within experimental error, the antigen binding site is identical between the original and fully modified IgGs. In all four structures, the quality of the electron density for BCMA is highest in the region of the binding site, particularly the segment containing the D*x*L loop. Aside from minor variations in sidechain conformation, this loop is positioned identically in the binding cavities of all antibody variants (Fig 3), with a nearly perfect superposition of L17, the critical residue recognized by all known BCMA binders, when the complete complex structures are aligned.

**Figure 4.**
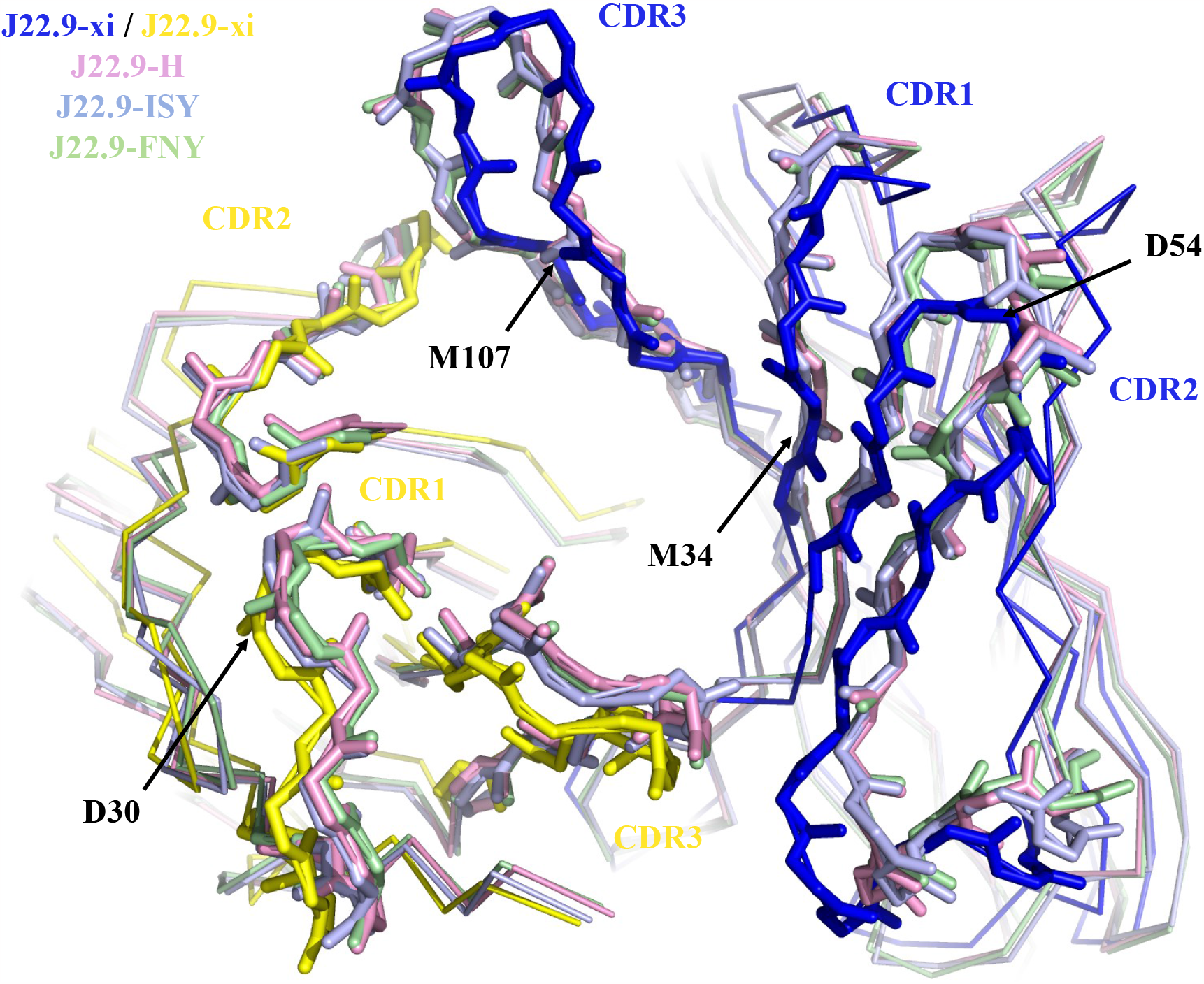
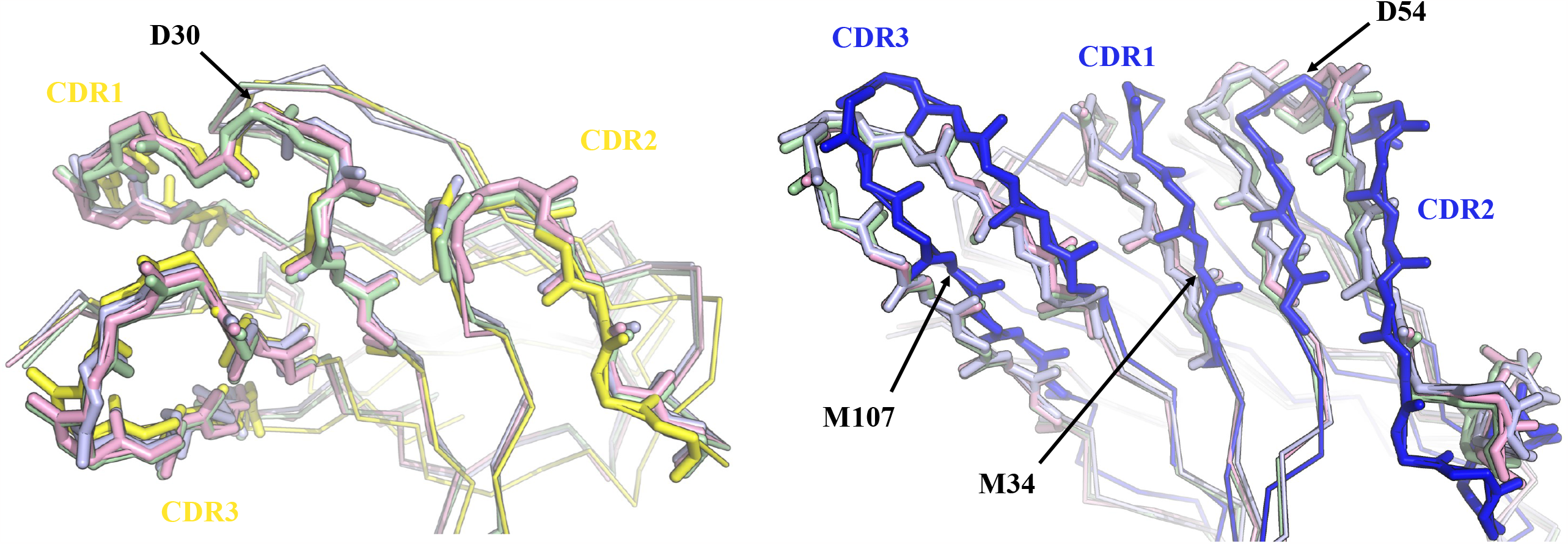
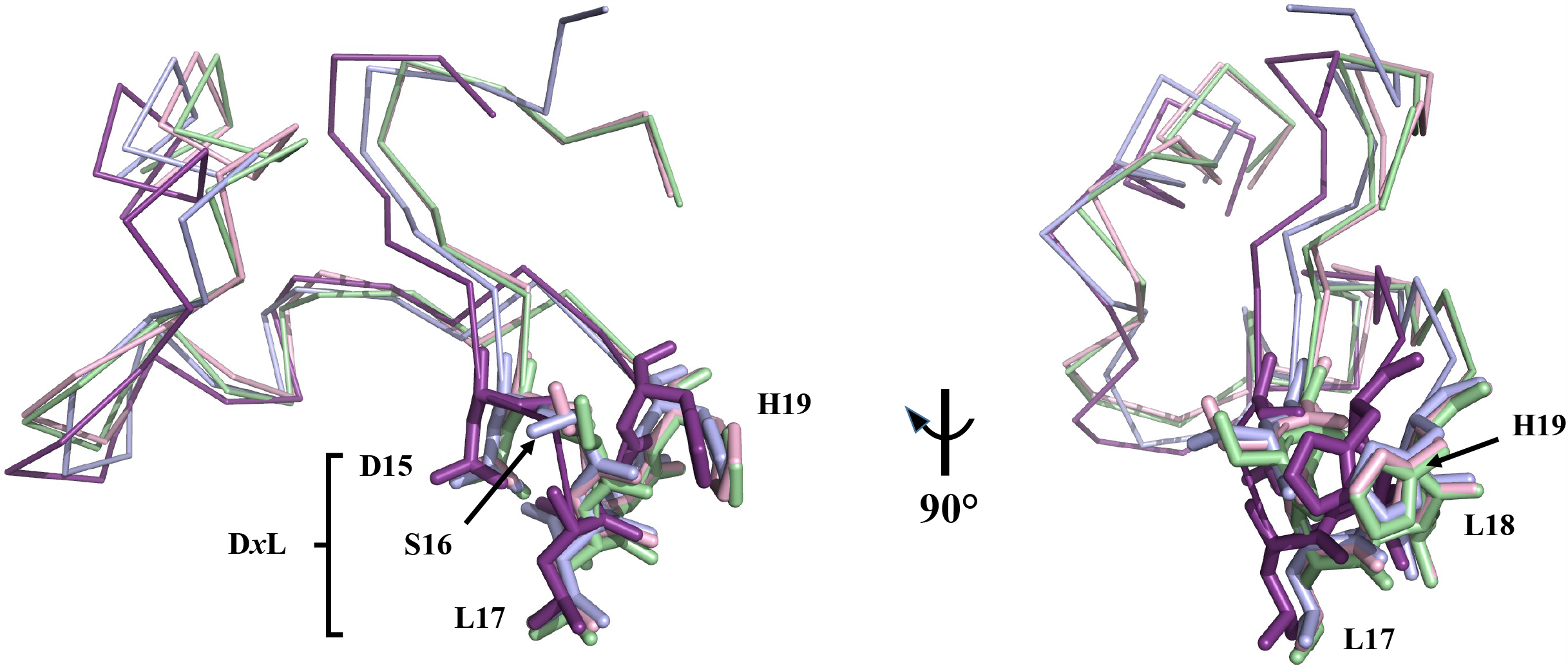
Structural alignments of J22.9-xi with humanized and optimized variants. A. View into the binding pocket of all four J22.9 aligned variants with BCMA removed. Only the backbone atoms are shown as lines, with the CDRs depicted as sticks. The V_H_ CDRs of J22.9-xi are colored blue and the V_L_ CDRs are colored yellow with J22.9-H shown in magenta, J22.9-ISY in light blue and J22.9-FNY in light green. The positions of all 4 PTM modifications are indicated. B. Separate views of the V_L_ (left panel) and V_H_ (right panel) domains with CDR depictions and coloring as in A. The conformations of all V_L_ CDR loops are nearly identical, as are those of the V_H_ CDRs of the humanized variants that all show a slight shift from the corresponding positions in J22.9-xi. C. Overlay of BCMA from all J22.9 variants generated by alignment of all 4 complete complex structures. BCMA bound to J22.9 -xi is shown in violet and BCMA molecules from the humanized structures are colored as per their corresponding V_L_ and V_H_ chains in A. The D*x*L loop residues are indicated.

The J22.9-FNY structure also shows retention of the important binding interactions between BCMA and IgG and only minor shifts in the backbone positions of the CDR loops despite the large NAG modification of V_H_ CDR2 (Fig. 5). Here again, the positions of the D*x*L loop and L17 are nearly identical to those seen in the J22.9-xi structure. That structure showed the minimal interaction of CDR2 with BCMA, allowing the possibility that substantial changes to this loop could be tolerated without eliminating ligand binding. Challenging this supposition with the glycosylation confirmed that productive changes in the CDR2 loop were possible, further emphasizing the value of the structure-based approach.

**Figure 5.**
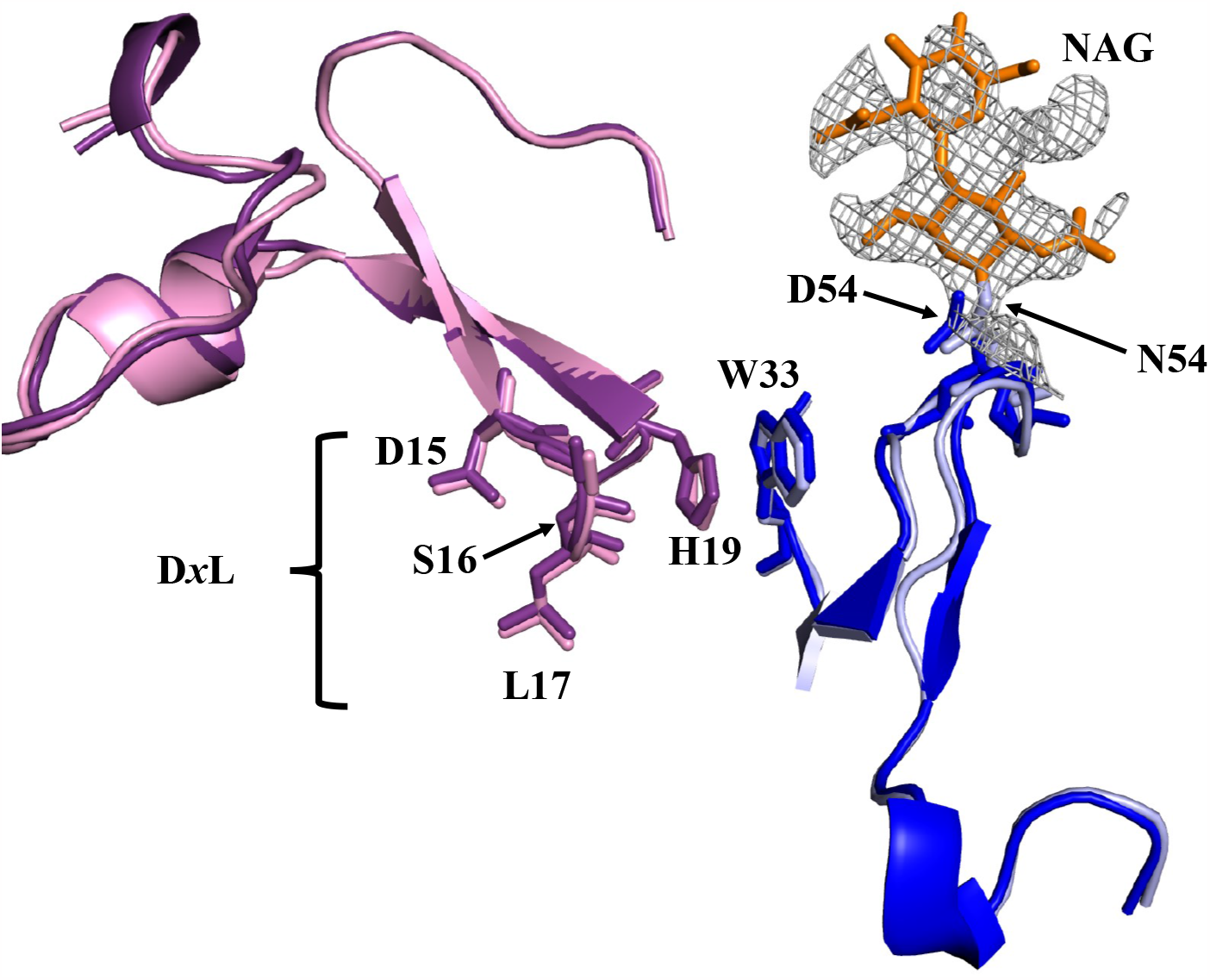
J22.9-H and J22.9-FNY binding site comparison. Structural alignment of J22.9-H:BCMA and J22.9-FNY:BCMA showing the interaction of BCMA with CDR2 and the position of the NAG modification.. The J22.9-H V_H_ is colored blue and the the J22.9-FNY V_H_ is colored light blue. The N-acetylglucosamine (NAG) modification on J22.9-FNY is colored orange. BCMA bound to J22.9-H is shown in violet and BCMA bound to J22.9 -FNY in magenta. The NAG modification in J22.9-FNY induces no substantial changes in the CDR2 loop. The most important features of both binding partners are labeled, including the D*x*L loop of BCMA, showing that the central L17 residue occupies a nearly identical position in both structures relative to the antibody. The single most extensive interaction in the complex, between W33 in V_H_ CDR1 and H19 in BCMA, is also explicitly shown and is unchanged between the two structures. The electron density (contoured at 1.5σ) for the NAG modification of J22.9-FNY is depicted as a grey mesh.

## Discussion

High resolution structural data is an invaluable resource for the characterization and development of therapeutic proteins. The successful, structure-based humanization of J22.9-xi, a tumor inhibiting IgG directed against MM, illustrates several of the advantages justifying the investment required to generate such data.

The humanization of J22.9-xi – a necessary step for evaluation of the molecule in the clinic – was greatly facilitated by the availability of the X-ray structure. Because each of the 41 amino acid substitutions indicated for complete conversion of the mouse to human variable domains could be vetted by building into the existing J22.9-xi structure, it was possible to flag potentially unsuitable changes before testing them in the lab. This allowed the identification of the A46 position that was then taken into account when planning the production of the humanized constructs. Using a cassette cloning strategy to rapidly evaluate groups of substitutions together, the humanizing A46L substitution identified by modeling was quickly isolated and verified to be the single one that could not be exchanged. Though the overall lack of destabilizing substitutions was fortuitous in this case, this cassette strategy is certainly applicable to more challenging molecules. Since the number and positions of cassette boundaries can be chosen as needed, problematic substitutions can be readily isolated, provided prior identification is possible.

The J22.9-xi structure also enabled evaluation of changes directly to the CDR loops for enhancing both the binding affinity for BCMA and the long-term stability of the IgG by targeting residues susceptible to PTM. The improvement in affinity validated the choices of replacement residues and the J22.9-ISY structure showed that, despite significant changes in their sequences, the conformations of the optimized CDRs did not substantially differ from those of the unmodified loops. The binding mode of BCMA in all cases was verified by the X ray structures to remain identical despite a total of 44 amino acid changes between J22.9-xi and the fully humanized/optimized variants, within experimental error. This successful optimization would also have been extremely difficult to plan and execute in the absence of the original structural data.

The structure further allowed testing of the flexibility of the V_H_ CDR2 by the generation of a glycosylation signal sequence. Using the PTM modification D54N, we induced an N-linked glycosylation of V_H_ CDR2 and solved the structure of this variant, J22.9-FNY, bound to BCMA. Comparison with the structure of the fully humanized J22.9-H again demonstrated the retention of the BCMA:IgG binding mode despite a large substitution in the immediate vicinity. These observations thereby indicate a site for further modification of the antibody, for example, the addition of detection labels/tracers, toxins, etc., that would have certainly remained undetected in the absence of a structure.

The most remarkable feature of the J22.9-xi:BCMA interaction was the minimal number of direct contacts between antibody and antigen, despite the picomolar affinity. Due to the high resolution of that structure, it was possible to identify many indirect contacts between the binding partners over “bridging” water molecules^29^. In addition, a total of 8 water molecules are completely buried in the interface between J22.9-xi and BCMA, with 6 filling the void in the binding pocket around the side chain of L17 from the D*x*L loop. The resolutions of the humanized and optimized variants are not sufficiently high to allow unambiguous identification of water molecules in the binding pocket. It is therefore not possible from this data to establish how many of these buried water positions may still be occupied in the humanized variant complexes. However, the similarities in conformations of the CDRs and the position of BCMA in all cases point to a high probability that these waters play an important role in the binding interaction in all variants.

We present a facilitated humanization procedure for therapeutic antibodies. The rapid design and execution of the cassette approach was guided by X-ray structural data on the ligand complex of the initial chimeric molecule. Subsequent interaction measurements and structures of the optimized variants showed an unexpectedly large improvement in binding affinity and retention of the binding mode, thus validating the procedure, which is extendable to any protein construct intended for clinical application. These results emphasize the fundamental importance of structural data for efficiently tailoring the properties of protein therapeutics. The success of this procedure has been independently validated for the humanized and optimized variant J22.9-ISY that retains its specificity for BCMA-positive tumors and, as an antibody-drug conjugate, exceeds its efficacy *in vivo*^30^.

## Acknowledgements

We are grateful to Susanne Scheu for expert technical assistance and acknowledge Felix Oden for FACS. We thank Thomas Tiller for fruitful discussion and for sharing many insights into antibody cloning and stability. We gratefully acknowledge the staff at BESSY Synchrotron at the Helmholtz Zentrum Berlin where all diffraction data was collected and in particular Manfred Weiss for access to and training on the HC-1 dehydration instrument. We additionally thank Katja Fälber for assistance with crystal screening and Luke Buchanan for critical reading of the manuscript. We are indebted to Mike Strauss for access to and assistance with his computing cluster at McGill University.

## References

1. Martin, K.P., Grimaldi, C., Grempler, R., Hansel, S. & Kumar, S. Trends in industrialization of biotherapeutics: a survey of product characteristics of 89 antibody-based therapeutics. MABS, 15, 2191301 (2023).

2. Lo K-M, Leger O, Hock B. Antibody engineering. Microbiol Spectrum 2(1):AID-0007-2012 (2014).

3. Morrison, S.L., Johnson, M.J., Herzenberg, L.A. & Ol V.T. Chimeric human antibody molecules: Mouse antigen-binding domains with human constant region domains. PNAS 81, 6851–6855 (1984).

4. Boulianne, G.L., Hozumi, N. & Shulman, M.J. Production of functional chimaeric mouse/human antibody. Nature, 312, 643–646 (1984).

5. Neuberger, M.S. et al. A hapten-specific chimaeric IgE antibody with human physiological effector function. Nature, 314, 268–270 (1985).

6. Jones, P.T., Dear, P.H., Foote, J., Neuberger, M.S. & Winter, G. Replacing the complementarity-determining regions in a human antibody with those from a mouse. Nature, 321, 522–525 (1986).

7. Kashmiri, S.V.S., DePascalis, R., Gonzales, N.R. & Schlom, J. SDR grafting – a new approach to antibody humanization. Methods, 36, 25–34 (2005).

8. Jespers, L.S., Roberts, A., Mahler, S.M., Winter, G. & Hoogenboom, H.R. Guiding the selection of human antibodies from phage display repertoires to a single epitope of an antigen. Biotechnology, 12, 899–903 (1994).

9. Osbourn, J., Groves, M. & Vaughan, T. From rodent reagents to human therapeutics using antibody guided selection. Methods, 36, 61–68 (2005).

10. Xing, L.X., Liu, Y. & Liu, J. Targeting BCMA in Multiple Myeloma: Advances in Antibody-Drug Conjugate Therapy. Cancers, 15, 2240–2255 (2023).

11. Berdeja, J.G. et al. Plain language summary of the CARITUDE-1 study of ciltacabtagene autoleucel for the treatment of people with relapsed or refractory multiple myeloma. Future Oncol., 19, 1235–1247 (2023).

12. Patiño-Escobar, B., Talbot, A. & Wiita, A.P. Overcoming proteasome inhibitor resistance in the immunotherapy era. Trends Pharmacol. Sci., 44, 507–518 (2023).

13. Darce, J.R., Arendt, B.K., Wu, X. & Jelinek, D.F. Regulated expression of BAFF-binding receptors during human B cell differentiation. J. Immunol., 179, 7276–7286 (2007).

14. Benson, M.J. et al. Cutting edge: the dependence of plasma cells and independence of memory cells on BAFF and APRIL. J. Immunol., 180, 3655–3659 (2008).

15. Good, K.L., Avery, D.T. & Tangye S.G. Resting human memory B cells are intrinsically programmed for enhanced survival and responsiveness to diverse stimuli compared to naïve B cells. J. Immunol., 182, 890–901 (2009).

16. Bluhm, J. et al. CAR T Cells with Enhanced Sensitivity to B Cell Maturation Antigen for the Targeting of B Cell Non-Hodgkin’s Lymphoma and Multiple Myeloma. Molecular Therapy, 26, 1906–1920 (2018).

17. Oden, F. et al. Potent anti-tumor response by targeting B cell maturation antigen (BCMA) in a mouse model of multiple myeloma. Molecular Oncology, 9, 1348–1358 (2015).

18. Emsley, P. & Cowtan, K. Coot: model-building tools for molecular graphics. Acta Crystallogr., D60, 2126–2132 (2004).

19. Sanchez-Weatherby, J. et al. Improving diffraction by humidity control: a novel device compatible with X-ray beamlines. Acta Cryst. D, 65, 1237–1246 (2009).

20. McCoy, A.J. et al. Phaser crystallographic software. J. Appl. Crystallogr., 40 (Pt 4), 658–674 (2007).

21. Adams, P.D. et al. PHENIX: a comprehensive Python-based system for macromolecular structure solution. Acta Cryst., D66, 213–221 (2010).

22. Krissinel E, Henrick K (2004). Secondary-structure matching (SSM), a new tool for fast protein structure alignment in three dimensions Acta Crystallogr. D60, 2256–2268

23. The PyMOL Molecular Graphics System, Version 1.2r3pre, Schrödinger, LLC

24. Ye, J., Ma, N., Madden, T.L. & Ostell, J.M. IgBLAST: an immunoglobulin variable domain sequence analysis tool. Nucleic Acids Research, 41, (Web Server Issue) W34–W40 (2013).

25. Alam, M.E. et al. Unique Impacts of Methionine Oxidation, Tryptophan Oxidation, and Asparagine Deamidation on Antibody Stability and Aggregation. J. Pharma. Sci., 109, 656–669 (2020).

26. Zheng, K. et al. Monoclonal Antibody Aggregation Associated with Free radical induced Oxidation. Int. J. Mol. Sci., 22, 3952–3967 (2021).

27. Vlasak, J. & Ionescu, R. Fragmentation of monoclonal antibodies. mAbs, 3, 253–263 (2011).

28. Joshi, A.B., Sawal, M., Kearney, W.R. & Kirsch, L.E. Studies on the Mechanism of Aspartic Acid Cleavage and Glutamine deamidation in the Acidic Degradation of Glucagon. J. Pharmaceut. Sci., 94, 1912–1927 (2005).

29. Marino, S.F., Olal, D. & Daumke, O. A complex water network contributes to high-affinity binding in an antibody-antigen interface. DiB, 6, 394–397 (2016).

30. Figueroa-Vasquez, V. et al. HDP-101, and Anti-BCMAAntibody-Drgug-Conjugate, Safely Delivers Amanitin to Induce Cell Death in Proliferating and Resting Multiple Myeloma Cells. Mol Cancer Therap., 20, 367–378 (2021).

